# Finding mutations in all the wrong places: Prevalence of knock-down resistance F1534S mutations among *Aedes albopictus (family: Culicidae, order: Diptera)* in North Carolina

**DOI:** 10.1101/2022.02.08.479572

**Authors:** Haley A. Abernathy, Brandon D. Hollingsworth, Dana A. Giandomenico, Kara A. Moser, Jonathan J. Juliano, Natalie M. Bowman, Phillip J. George, Michael H. Reiskind, Ross M. Boyce

## Abstract

**Background:** Knock-down resistance (*kdr*) mutations in the voltage gated sodium channel gene of *Aedes* species mosquitoes are biomarkers for resistance to pyrethroid insecticides. In the United States, few studies have reported *kdr* mutations among *Aedes albopictus* (*family: Culicidae, order: Diptera, Skuse, 1895)* populations. In this study we sought to explore the potential for permethrin-impregnated uniforms worn by military servicemembers to drive *kdr* emergence.

**Methods and Results:** We collected 538 *Aedes albopictus* mosquitoes, including 156 from 4 sites at Fort Bragg (exposed), North Carolina and 382 from 15 sites in Wake County (control), North Carolina to compare the prevalence of *kdr* mutations. Of those successfully sequenced, we identified 12 (3.0%) mosquitoes with *kdr* mutations, all of which were attributed to variants at position 1534 within domain 3. All mutations were found in mosquitoes collected at Wake County sites; no mutations were identified at Fort Bragg. There was a focus of mutations observed at the Wake County sites with approximately 92% (11 of 12) of the mosquitoes with the mutation coming from one site, where *kdr* mutations represented 24.4% (11 of 45) of all mosquitoes collected.

**Conclusions:** Our study did not show any evidence that universal implementation of permethrin-impregnated uniforms drives the development of resistance. In contrast, we observed highly focal resistance in a suburban area of Raleigh, which may be attributable to peri-domestic mosquito control activities that involve area dispersal of pyrethroid insecticides. More robust surveillance is needed to monitor the emergence and spread of resistance.

**AUTHOR SUMMARY:** Resistance to commonly employed insecticides among *Aedes albopictus* (*family: Culicidae, order: Diptera, Skuse, 1895)* mosquitoes poses as a substantial public health threat. In this study we sought to explore the potential for permethrin-impregnated uniforms worn by military servicemembers to drive emergence of resistance to pyrethroid insecticides by collecting and testing mosquitoes from both military and civilian sites. Overall, we did not identify mosquitoes harboring resistance at Ft. Bragg, but did find a focus of resistance in sub-urban Raleigh, which may be driven by commercial, peri-domestic mosquito control activities. These results suggest that resistance to pyrethroid insecticides may be more prevalent in the United States than previously known, but highly heterogenous. More robust surveillance is needed to monitor the emergence and spread of resistance.

## INTRODUCTION

*Aedes albopictus* (*family: Culicidae, order: Diptera, Skuse, 1895)* is a highly invasive mosquito species and competent vector of several pathogens of public health importance including West Nile virus, eastern equine encephalitis virus, and LaCrosse virus, the most common cause of pediatric encephalitis in the United States (Westby et al. 2015, Vanlandingham et al. 2016, Gloria-Soria et al. 2020). Though not the most important vector of major arboviral pathogens such as dengue and chikungunya, *Ae. albopictus* has been incriminated in several large outbreaks (Effler et al. 2005, Gérardin et al. 2008). In the United States, *Ae. albopictus* is more common than *Aedes aegypti*, often reaching higher population densities and placing them in an optimal position to transmit human and animal disease (Khan et al. 2020). In addition, they are known to be aggressive biters, making them a major nuisance concern.

Organophosphate and pyrethroid-based insecticides have been the primary method of controlling adult mosquito populations in the United States, with pyrethroids being the more widely accepted, less toxic, and cost-effective tool in the last 20 years (Estep et al. 2018, Stoops et al. 2019). Commercially available pyrethroids like permethrin, deltamethrin, and bifenthrin, are widely utilized for *Ae. albopictus* control. Although not widely reported, resistance to pyrethroids among *Ae. albopictus* has been documented in the United States (Marcombe et al. 2014, Xu et al. 2016, McInnis et al. 2019, Richards et al. 2019). The most common pyrethroid resistance mechanism, knockdown resistance (*kdr*), affects the voltage-gated sodium channel (VGSC) target site of pyrethroid insecticides. There are several VGSC gene mutations that serve as biomarkers for pyrethroid resistance in *Ae. albopictus*. The most commonly found single nucleotide polymorphism (SNP) conferring insecticide resistance occurs at position 1016, where valine is replaced with isoleucine or glycine (V1016I or V1016G) (Auteri et al. 2018). Other important SNPs include replacement of isoleucine with threonine at position 1532 (I1532T) and replacement of phenylalanine with serine, leucine, or cysteine at position 1534 (F1534S, F1534L, F1534C).

The emergence of pyrethroid resistance in *Ae. albopictus* represents a major threat to global public health.^16^ The widespread use of pyrethroids has catalyzed the rapid evolution of resistance to pyrethroids among mosquitoes in several countries (Auteri et al. 2018). For example, in the city of Guangzhou, China, vector control measures were implemented following a surge of dengue cases in 2014. Incident rates decreased in the following years but quadrupled from 2018 to 2019 [17]. This increase in cases was preceded by several reports of increased pyrethroid resistance caused by F1534S and F1534L mutations in *Ae. albopictus* populations in Guangzhou (Li et al. 2018, Su et al. 2019).

Mosquito control is heterogeneously employed in the United States, with some states having very active control districts. In North Carolina in particular, control measures vary substantially between counties (Del Rosario et al. 2014). Private, peri-domestic focused mosquito control accounts for a significant proportion of mosquito control in North Carolina but is highly variable with regards to region and socioeconomic status. In addition, among select populations, such as military service members, there exists a nearly universal usage of permethrin-impregnated uniforms designed to protect against vector-borne diseases both at home and when deployed abroad (Armed Forces Pest Management Board 2015). Widespread use of these uniforms potentially creates selective pressure for the development of pyrethroid resistance, especially in and around major military installations. Given near universal use of permethrin-impregnated uniforms at Fort Bragg, the largest US military installation by population, we hypothesized that pyrethroid resistance may have emerged.

The overarching goal of this study was to compare the prevalence of *kdr*-mutations among mosquitoes collected at Fort Bragg to mosquitoes collected in the Raleigh metropolitan area in Wake County, NC, where permethrin-treated clothing is less commonly used. To test this hypothesis, we tested individual *Ae. albopictus* collected across sites at Fort Bragg and in Wake County using sequence analysis targeting known mutations associated with *kdr*.

## METHODS

### Mosquito Collections

In August and September of 2020, we collected adult *Ae. albopictus* mosquitoes from 4 sentinel surveillance sites in Fort Bragg, NC over 15 total trap-nights using CDC light traps. Similarly, we collected *Ae. albopictus* mosquitoes from 15 sentinel surveillance sites in Wake County, NC over 15 total trap-nights using CDC light traps. Mosquitoes were identified to species using standard keys and were grouped by sex, species, and collection site before being stored at -20° C in 95% ethanol until extraction (Harrison BA. et al. 2016) Collection data for Wake County and Fort Bragg can be found in Additional Data 1: Dataset S1 and S2 respectively.

### Extraction and Sequencing

DNA from individual whole mosquitoes was extracted using the Quick-DNA Tissue/Insect Miniprep Kit (Zymo Research, Irvine, CA) per the manufacturer protocol. Partial sequences from domain 2 and 3 of the VGSC gene, encompassing amino acid residue positions 1016, 1532 and 1534, were amplified using previously published primer sets and the FastStart™ High Fidelity PCR kit (Bowman et al. 2018, Zhou et al. 2019). Primer sequences are shown in **Table 1**. Reactions for both domains were run based on a previously published qPCR protocol with several modifications: 17.25 uL water, 2.5 uL of template DNA, 1 uL of forward primer, and 1 uL of reverse primer (Bowman et al. 2018). Primers K21 and V3R were used to sequence domain 2 and domain 3 respectively. Sequences (50 base-pairs in length) for domain 2 and domain 3 encompassing the positions of interest were aligned and analyzed using the Geneious Prime software (Biomatters, Auckland, New Zealand). Individual mosquito sequence data for domain two and domain three can be found in Additional Data 1: Dataset S3 and S4 respectively.

**Table 1:**
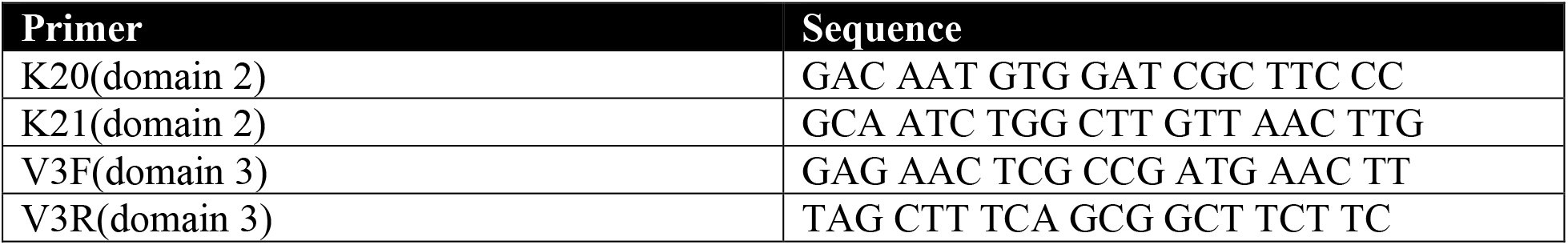
Primer sequences used to partially amplify domain 2 and domain 3 of the voltage-gated sodium channel gene in *Ae. albopictus*.

### Data Analysis

We recorded the number of *Ae. albopictus* mosquitoes collected, stratified by site of collection. Collection sites within Wake County were considered residential if surrounded by buildings within 100 meters on all sides. Comparisons between locations were done using a difference in proportions test. In addition, individual sites were tested against the location mean to identify sites with higher levels of resistance. All analysis was performed using R Statistical Software and surveillance sites were depicted on a map using the OpenStreetMap package (R Core Team 2018, Fellows 2019)

## RESULTS

From August to September 2020, we collected 538 (28 males and 510 females) *Ae. albopictus*. 156 were collected from 4 sites at Fort Bragg and 382 from 15 sites in Wake County. Domain 2 of the VGSC gene was successfully sequenced in 467 (86.8%) mosquitoes, with similar rates between Fort Bragg (129, 82.7%) and Wake County (338, 88.5%, p=0.07). Domain 3 of the VGSC gene was successfully sequenced in 412 (76.6%) mosquitoes, with modestly higher success rates among mosquitoes from Fort Bragg (134, 85.9%) compared to those from Wake County (278, 72.8%, p=.001).

12 *Ae. albopictus,* 2 males and 10 females, with *kdr* mutations were identified, all of which were variants at position 1534 in domain 3 with serine replacing phenylalanine. All mutations were identified in *Ae. albopictus* collected at Wake County sites (**Table 2**); no mutations were identified in mosquitoes collected from Fort Bragg. The prevalence of *kdr* mutations among sequenced mosquitoes from the Wake County sites was 4.3% (12 of 278). There was an apparent focus of mutations observed at the Wake County sites with 91.6% (11 of 12) coming from one site (**Table 3**). At this site, *Ae. albopictus* harboring *kdr* mutations represented 24.4% (11 of 45) of all mosquitoes collected. This was significantly different than what was seen in Wake County as a whole (p < .001). A visual representation of Wake County and Fort Bragg collection sites, the number of mosquitoes sampled from each site, and the *kdr* mutation prevalence for each site can be found in **Figure 1**.

**Table 2:** Sequencing results for *kdr* mutations stratified by collection site and domain.

|  | Domain 2 |  | Domain 3 |  |
| --- | --- | --- | --- | --- |
| Location | Wild Type | Mutant | Wild Type | Mutant |
| Fort Bragg | 129 | 0 | 134 | 0 |
| Wake County | 338 | 0 | 266 | 12 |

**Table 3:** Domain 3 *kdr* mutation counts found in Wake County separated into sites.

| Location | Site | Total Sequenced | *Kdr* mutations |
| --- | --- | --- | --- |
| Wake County | Mission | 19 | 0 (0.0%) |
| Wake County | Trexler | 24 | 0 (0.0%) |
| Wake County | Varsity | 11 | 0 (0.0%) |
| Wake County | VDR | 45 | 11 (24.4%) |
| Wake County | BRF | 31 | 1 (3.2 %) |
| Wake County | Clarion | 38 | 0 (0.0%) |
| Wake County | Dairy | 69 | 0 (0.0%) |
| Wake County | Farm | 18 | 0 (0.0%) |
| Wake County | Other | 23 | 0 (0.0%) |
| Fort Bragg | Woodland Heights | 45 | 0 (0.0%) |
| Fort Bragg | Ryders Golf Course | 25 | 0 (0/0%) |
| Fort Bragg | Nijmegen | 60 | 0 (0/0%) |
| Fort Bragg | Ritz-Epps | 4 | 0 (0/0%) |

**Figure 1:**
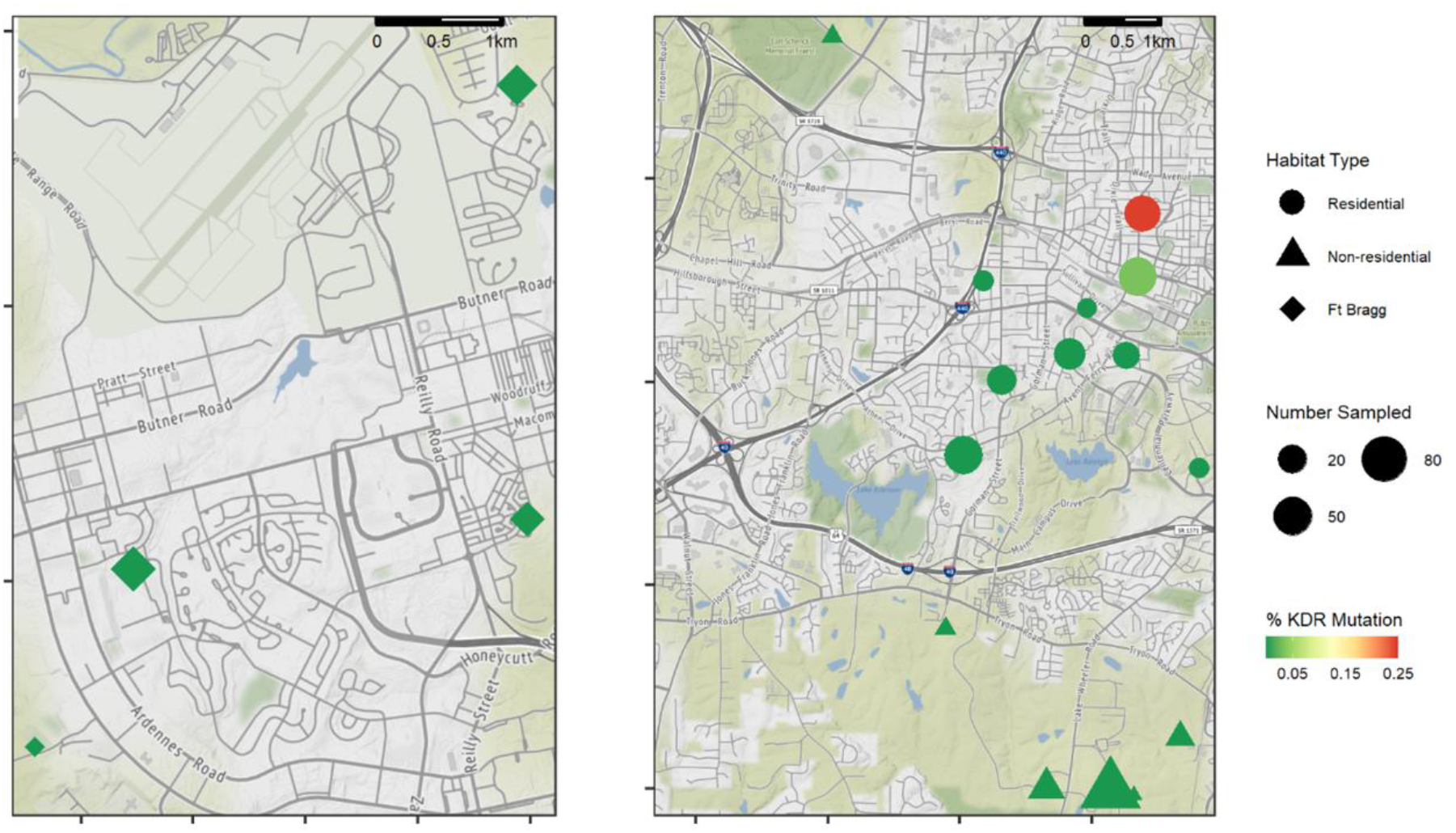
Map of collection sites at Fort Bragg (left) and Wake County (right) showing mosquito collection sites, number of *Ae. albopictus* mosquitoes collected, and prevalence of *kdr* mutations among collected mosquitoes.

## DISCUSSION

Contrary to our hypothesis that widespread use of permethrin-impregnated uniforms might drive resistance, we detected no *kdr* mutations in the 156 *Ae. albopictus* collected from Fort Bragg. In contrast, we identified a high prevalence of *kdr* mutations in mosquitoes collected from a single residential site in Wake County, North Carolina. These findings suggest that foci of resistant mosquitoes may exist, with peri-domestic mosquito abatement (e.g., private backyard control) activities being a potential driver. While our results cannot define the geographic extent of resistance, they clearly support the need for increased insecticide resistance surveillance efforts in suburban and semi-urban areas.

Our finding that nearly 25% of *Ae. albopictus* at a single site harbor the *kdr* mutation is important because *Ae. albopictus* are capable of transmitting several diseases already present in North Carolina (e.g., West Nile virus and LaCrosse encephalitis). Additionally, *Ae. albopictus* would likely be the primary vector of other arboviral diseases if introduced into North Carolina. They also play a large role in the transmission of dog heartworms, a nationwide problem with approximately one million dogs affected each year (Spence Beaulieu and Reiskind 2020). Widespread prevalence of *kdr* mutations could severely limit the ability to control and respond to outbreaks of these diseases, as well as impact the effectiveness of permethrin-treated uniforms as originally hypothesized (Estep et al. 2020).

The heterogenous distribution of resistance found in this study is consistent with previously published work on *kdr* mutations in *Ae. albopictus* (Xu et al. 2016). The high rate of *kdr* mutations seen at one wake county site may be a result of peri-domestic, commercial abatement employed to reduce *Ae. albopictus* nuisance biting. In contrast, the absence of *kdr* mutations at Fort Bragg could be due to a number of causes. Permethrin-impregnated uniforms substantially reduce landing and biting rates as they both repel and kill mosquitoes (Orsborne et al. 2016). This combination may not provide sufficient selective pressure as the widespread spraying of insecticide. In addition, Fort Bragg does not currently carry out any large-scale mosquito abatement measures suggesting a lack of selective pressure due to the absence of widespread insecticide use on base. Lastly, the sample size and number of collection sites at Fort Bragg were much smaller in comparison to Wake County. It is possible that *kdr* mutations may exist at Fort Bragg but were not discovered by our study due to the more limited collection.

Our study had several limitations including a relatively short collection period, the use of already established collection sites not specifically chosen for this analysis, and the assumption that resistance exists among *kdr* mutant mosquitoes. Additionally, mosquito collection efforts were less extensive at Fort Bragg when compared to Wake County. Our study also had several strengths including the quasi-experimental design with intervention (Fort Bragg) and control (Wake County) areas as well as the use of well-established sequencing methods.

## CONCLUSION

Though resistance is not widely reported in the United States, our study demonstrates that *kdr* mutations may be common. Our findings suggest that more robust surveillance is needed to monitor trends in resistance and define appropriate insecticides for disease control as part of integrated pest management strategies. Future studies need to more comprehensively gauge the extent of mutations among *Ae. albopictus* in North Carolina and determine underlying causes of mutations beyond being located in residential areas.

## ABBREVIATIONS

VGSC: voltage-gated sodium channel
*kdr*: knock-down resistance.

## SUPPLEMENTARY INFORMATION

**Additional file 1: Table S1.** Mosquito collection data from Wake County. **Table S2.** Mosquito collection data from Fort Bragg. **Table S3.** Domain two sequence data for each mosquito successfully sequenced. **Table S4.** Domain three sequence data for each mosquito successfully sequenced.

## DECLARATIONS

## Previous Publication

The authors confirm that the submitted work has not been previously presented or published in any format and is not under consideration at any other journal.

## Author Contributions

Study conception and design: RMB, MHR, CM, PG, NMB. Funding: RMB, NMB, CM, MHR, PG. Study implementation: RMB, MHR, PG, CM, HAA, NMB, JJJ. Data analysis: HA, BDH, RMB. First draft of manuscript: HA, BDH, DAG, RMB. Revisions: All.

## Availability of Data and Materials

Deidentified individual data that supports the results will be shared beginning 9 to 36 months following publication provided the investigator who proposes to use the data has approval from an Institutional Review Board (IRB), Independent Ethics Committee (IEC), or Research Ethics Board (REB), as applicable, and executes a data use/sharing agreement with UNC.

## Acknowledgements

None

