## Supplemental Tables for "Finding mutations in all the wrong places: Prevalence of knock-down resistance F1534S mutations among *Aedes albopictus (family: Culicidae, order: Diptera)* in North Carolina"

### SUPPLEMENTARY DATA

**Table S1.** Mosquito collection data from Wake County.

| Site ID | Date Set | Set Time | Date Pickup | Pickup Time | Female | Male |
|---|---|---|---|---|---|---|
| DIX | 8/26/2020 | 1245 | 8/27/2020 | 0950 | 2 | |
| Schenck | 8/27/2020 | 1240 | 8/28/2020 | 1045 | 2 | 1 |
| Grinnells | 8/26/2020 | 1210 | 8/27/2020 | 0915 | 4 | |
| MET | 8/27/2020 | 1325 | 8/28/2020 | 0820 | 6 | |
| Market | 8/25/2020 | 1240 | 8/26/2020 | 0920 | 8 | |
| Forensic | 8/25/2020 | 1325 | 8/26/2020 | 0950 | 11 | |
| Farm | 8/26/2020 | 1315 | 8/27/2020 | 1050 | 20 | |
| Mission | 8/25/2020 | 1210 | 8/26/2020 | 0900 | 22 | |
| Trexler | 8/27/2020 | 1340 | 8/28/2020 | 0810 | 24 | 1 |
| Varsity | 8/26/2020 | 1220 | 8/27/2020 | 0925 | 32 | |
| VDR | 8/27/2020 | 1225 | 8/28/2020 | 1030 | 42 | 3 |
| BRF | 8/27/2020 | 1215 | 8/28/2020 | 1025 | 45 | 9 |
| Clarion | 8/26/2020 | 1230 | 8/27/2020 | 0935 | 49 | 11 |
| Dairy | 8/26/2020 | 1330 | 8/27/2020 | 1100 | 87 | 2 |
| Office | 8/25/2020 | 1315 | 8/26/2020 | 0940 | | 1 |
| | | | | **SUM** | 354 | 28 |

**Table S2.** Mosquito collection data from Fort Bragg.

| Site ID | Date Set | Time Set | Date Pickup | Pickup Time | Females | Males |
|---|---|---|---|---|---|---|
| Nijmegen | 9/10/2020 | 14:13 | 9/11/2020 | 14:12 | 26 | |
| Nijmegen | 8/31/2020 | 14:50 | 9/1/2020 | 15:13 | 2 | |
| Nijmegen | 9/2/2020 | 14:11 | 9/3/2020 | 13:48 | 23 | |
| Nijmegen | 9/8/2020 | 14:08 | 9/9/2020 | 14:45 | 17 | |
| Ritz-Epps | 8/31/2020 | 15:03 | 9/1/2020 | 15:05 | 1 | |
| Ritz-Epps | 9/2/2020 | 14:25 | 9/3/2020 | 14:01 | 2 | |
| Ritz-Epps | 9/8/2020 | 14:22 | 9/9/2020 | 14:20 | 2 | |
| Ryder Golf Course | 9/2/2020 | 14:53 | 9/3/2020 | 13:22 | 5 | |
| Ryder Golf Course | 9/8/2020 | 13:53 | 9/9/2020 | 15:23 | 8 | |
| Ryder Golf Course | 9/10/2020 | 14:53 | 9/11/2020 | 13:37 | 14 | |
| Ryder Golf Course | 8/31/2020 | 14:35 | 9/1/2020 | 15:44 | 6 | |
| Woodland Heights | 8/31/2020 | 15:35 | 9/1/2020 | 15:38 | 15 | |
| Woodland Heights | 9/2/2020 | 13:55 | 9/3/2020 | 13:35 | 19 | |
| Woodland Heights | 9/8/2020 | 13:33 | 9/9/2020 | 15:00 | 1 | |
| Woodland Heights | 9/10/2020 | 13:54 | 9/11/2020 | 13:55 | 15 | |
| | | | | SUM | 156 | |

**Table S3.** Domain two sequence data for each mosquito successfully sequenced.

| Location | Position 1016 | KDR Mutant? | Site |
|---|---|---|---|
| Fort Bragg | GTG | no | Woodland Heights |
| Fort Bragg | GTG | no | Woodland Heights |
| Fort Bragg | GTT | no | Ryder Golf Course |
| Fort Bragg | GTT | no | Ryder Golf Course |
| Fort Bragg | GTT | no | Ryder Golf Course |
| Fort Bragg | GTG | no | Nijmegen |
| Wake County | GTG | no | Farm |
| Wake County | GTG | no | Dairy |
| Wake County | GTG | no | Forensic |
| Wake County | GTT | no | VDR |
| Wake County | GTA | no | Mission |
| Fort Bragg | GTA | no | Ritz-Epps |
| Fort Bragg | GTA | no | Woodland Heights |
| Fort Bragg | GTA | no | Woodland Heights |
| Fort Bragg | GTA | no | Woodland Heights |
| Fort Bragg | GTA | no | Woodland Heights |
| Fort Bragg | GTA | no | Woodland Heights |
| Fort Bragg | GTA | no | Woodland Heights |
| Fort Bragg | GTA | no | Woodland Heights |
| Fort Bragg | GTA | no | Woodland Heights |
| Fort Bragg | GTA | no | Woodland Heights |
| Fort Bragg | GTA | no | Woodland Heights |
| Fort Bragg | GTA | no | Woodland Heights |
| Fort Bragg | GTA | no | Woodland Heights |

| Location | Position 1016 | KDR Mutant? | Site |
|---|---|---|---|
| Fort Bragg | GTA | no | Woodland Heights |
| Fort Bragg | GTA | no | Ryder Golf Course |
| Fort Bragg | GTA | no | Ryder Golf Course |
| Fort Bragg | GTA | no | Ryder Golf Course |
| Fort Bragg | GTA | no | Ryder Golf Course |
| Fort Bragg | GTA | no | Nijmegen |
| Fort Bragg | GTA | no | Woodland Heights |
| Fort Bragg | GTA | no | Woodland Heights |
| Fort Bragg | GTA | no | Woodland Heights |
| Fort Bragg | GTA | no | Woodland Heights |
| Fort Bragg | GTA | no | Woodland Heights |
| Fort Bragg | GTA | no | Woodland Heights |
| Fort Bragg | GTA | no | Woodland Heights |
| Fort Bragg | GTA | no | Woodland Heights |
| Fort Bragg | GTA | no | Woodland Heights |
| Fort Bragg | GTA | no | Woodland Heights |
| Fort Bragg | GTA | no | Woodland Heights |
| Fort Bragg | GTA | no | Woodland Heights |
| Fort Bragg | GTA | no | Woodland Heights |
| Fort Bragg | GTA | no | Woodland Heights |
| Fort Bragg | GTA | no | Woodland Heights |
| Fort Bragg | GTA | no | Woodland Heights |
| Fort Bragg | GTA | no | Woodland Heights |
| Fort Bragg | GTA | no | Woodland Heights |
| Fort Bragg | GTA | no | Nijmegen |
| Fort Bragg | GTA | no | Nijmegen |

| Location | Position 1016 | KDR Mutant? | Site |
|---|---|---|---|
| Fort Bragg | GTA | no | Nijmegen |
| Fort Bragg | GTA | no | Nijmegen |
| Fort Bragg | GTA | no | Nijmegen |
| Fort Bragg | GTA | no | Nijmegen |
| Fort Bragg | GTA | no | Nijmegen |
| Fort Bragg | GTA | no | Nijmegen |
| Fort Bragg | GTA | no | Nijmegen |
| Fort Bragg | GTA | no | Nijmegen |
| Fort Bragg | GTA | no | Nijmegen |
| Fort Bragg | GTA | no | Nijmegen |
| Fort Bragg | GTA | no | Nijmegen |
| Fort Bragg | GTA | no | Nijmegen |
| Fort Bragg | GTA | no | Nijmegen |
| Fort Bragg | GTA | no | Nijmegen |
| Fort Bragg | GTA | no | Nijmegen |
| Fort Bragg | GTA | no | Nijmegen |
| Fort Bragg | GTA | no | Nijmegen |
| Fort Bragg | GTA | no | Ryder Golf Course |
| Fort Bragg | GTA | no | Ryder Golf Course |
| Fort Bragg | GTA | no | Ryder Golf Course |
| Fort Bragg | GTA | no | Ryder Golf Course |
| Fort Bragg | GTA | no | Ritz-Epps |
| Fort Bragg | GTA | no | Ryder Golf Course |
| Fort Bragg | GTA | no | Ryder Golf Course |
| Fort Bragg | GTA | no | Ryder Golf Course |
| Fort Bragg | GTA | no | Ryder Golf Course |

| Location | Position 1016 | KDR Mutant? | Site |
|---|---|---|---|
| Fort Bragg | GTA | no | Ryder Golf Course |
| Fort Bragg | GTA | no | Nijmegen |
| Fort Bragg | GTA | no | Nijmegen |
| Fort Bragg | GTA | no | Nijmegen |
| Fort Bragg | GTA | no | Nijmegen |
| Fort Bragg | GTA | no | Nijmegen |
| Fort Bragg | GTA | no | Nijmegen |
| Fort Bragg | GTA | no | Nijmegen |
| Fort Bragg | GTA | no | Nijmegen |
| Fort Bragg | GTA | no | Nijmegen |
| Fort Bragg | GTA | no | Nijmegen |
| Fort Bragg | GTA | no | Nijmegen |
| Fort Bragg | GTA | no | Ritz-Epps |
| Fort Bragg | GTA | no | Ritz-Epps |
| Fort Bragg | GTA | no | Woodland Heights |
| Fort Bragg | GTA | no | Woodland Heights |
| Fort Bragg | GTA | no | Woodland Heights |
| Fort Bragg | GTA | no | Woodland Heights |
| Fort Bragg | GTA | no | Woodland Heights |
| Fort Bragg | GTA | no | Woodland Heights |
| Fort Bragg | GTA | no | Woodland Heights |
| Fort Bragg | GTA | no | Woodland Heights |
| Fort Bragg | GTA | no | Woodland Heights |
| Fort Bragg | GTA | no | Ryder Golf Course |
| Fort Bragg | GTA | no | Ryder Golf Course |
| Fort Bragg | GTA | no | Ryder Golf Course |

| Location | Position 1016 | KDR Mutant? | Site |
|---|---|---|---|
| Fort Bragg | GTA | no | Ryder Golf Course |
| Fort Bragg | GTA | no | Ryder Golf Course |
| Fort Bragg | GTA | no | Ryder Golf Course |
| Fort Bragg | GTA | no | Ryder Golf Course |
| Fort Bragg | GTA | no | Ryder Golf Course |
| Fort Bragg | GTA | no | Ryder Golf Course |
| Fort Bragg | GTA | no | Ryder Golf Course |
| Fort Bragg | GTA | no | Ryder Golf Course |
| Fort Bragg | GTA | no | Ryder Golf Course |
| Fort Bragg | GTA | no | Nijmegen |
| Fort Bragg | GTA | no | Nijmegen |
| Fort Bragg | GTA | no | Nijmegen |
| Fort Bragg | GTA | no | Nijmegen |
| Fort Bragg | GTA | no | Nijmegen |
| Fort Bragg | GTA | no | Nijmegen |
| Fort Bragg | GTA | no | Nijmegen |
| Fort Bragg | GTA | no | Nijmegen |
| Fort Bragg | GTA | no | Nijmegen |
| Fort Bragg | GTA | no | Nijmegen |
| Fort Bragg | GTA | no | Nijmegen |
| Fort Bragg | GTA | no | Nijmegen |
| Fort Bragg | GTA | no | Nijmegen |
| Fort Bragg | GTA | no | Nijmegen |
| Fort Bragg | GTA | no | Nijmegen |
| Fort Bragg | GTA | no | Nijmegen |
| Fort Bragg | GTA | no | Nijmegen |

| Location | Position 1016 | KDR Mutant? | Site |
|---|---|---|---|
| Fort Bragg | GTA | no | Nijmegen |
| Fort Bragg | GTA | no | Nijmegen |
| Fort Bragg | GTA | no | Nijmegen |
| Fort Bragg | GTA | no | Nijmegen |
| Fort Bragg | GTA | no | Nijmegen |
| Fort Bragg | GTA | no | Nijmegen |
| Wake County | GTA | no | Mission |
| Wake County | GTA | no | Mission |
| Wake County | GTA | no | Mission |
| Wake County | GTA | no | Mission |
| Wake County | GTA | no | Mission |
| Wake County | GTA | no | Mission |
| Wake County | GTA | no | Mission |
| Wake County | GTA | no | Mission |
| Wake County | GTA | no | Mission |
| Wake County | GTA | no | Mission |
| Wake County | GTA | no | Mission |
| Wake County | GTA | no | Mission |
| Wake County | GTA | no | Mission |
| Wake County | GTA | no | Mission |
| Wake County | GTA | no | Mission |
| Wake County | GTA | no | Mission |
| Wake County | GTA | no | Market |
| Wake County | GTA | no | Market |
| Wake County | GTA | no | Market |
| Wake County | GTA | no | Market |

| Location | Position 1016 | KDR Mutant? | Site |
|---|---|---|---|
| Wake County | GTA | no | Office |
| Wake County | GTA | no | Forensic |
| Wake County | GTA | no | Forensic |
| Wake County | GTA | no | Forensic |
| Wake County | GTA | no | Forensic |
| Wake County | GTA | no | Forensic |
| Wake County | GTA | no | Forensic |
| Wake County | GTA | no | Forensic |
| Wake County | GTA | no | Grinnells |
| Wake County | GTA | no | Grinnells |
| Wake County | GTA | no | Grinnells |
| Wake County | GTA | no | Grinnells |
| Wake County | GTA | no | Varsity |
| Wake County | GTA | no | Varsity |
| Wake County | GTA | no | Varsity |
| Wake County | GTA | no | Varsity |
| Wake County | GTA | no | Varsity |
| Wake County | GTA | no | Varsity |
| Wake County | GTA | no | Varsity |
| Wake County | GTA | no | Varsity |
| Wake County | GTA | no | Varsity |
| Wake County | GTA | no | Varsity |
| Wake County | GTA | no | Varsity |
| Wake County | GTA | no | Varsity |
| Wake County | GTA | no | Varsity |
| Wake County | GTA | no | Varsity |

| Location | Position 1016 | KDR Mutant? | Site |
|---|---|---|---|
| Wake County | GTA | no | Varsity |
| Wake County | GTA | no | Varsity |
| Wake County | GTA | no | Varsity |
| Wake County | GTA | no | Varsity |
| Wake County | GTA | no | Varsity |
| Wake County | GTA | no | Varsity |
| Wake County | GTA | no | Varsity |
| Wake County | GTA | no | Varsity |
| Wake County | GTA | no | Varsity |
| Wake County | GTA | no | Varsity |
| Wake County | GTA | no | Varsity |
| Wake County | GTA | no | Varsity |
| Wake County | GTA | no | Varsity |
| Wake County | GTA | no | Varsity |
| Wake County | GTA | no | Varsity |
| Wake County | GTA | no | Varsity |
| Wake County | GTA | no | Varsity |
| Wake County | GTA | no | Clarion |
| Wake County | GTA | no | Clarion |
| Wake County | GTA | no | Clarion |
| Wake County | GTA | no | Clarion |
| Wake County | GTA | no | Clarion |
| Wake County | GTA | no | Clarion |
| Wake County | GTA | no | Clarion |
| Wake County | GTA | no | Clarion |
| Wake County | GTA | no | Clarion |

| Location | Position 1016 | KDR Mutant? | Site |
|---|---|---|---|
| Wake County | GTA | no | Clarion |
| Wake County | GTA | no | Clarion |
| Wake County | GTA | no | Clarion |
| Wake County | GTA | no | Clarion |
| Wake County | GTA | no | Clarion |
| Wake County | GTA | no | Clarion |
| Wake County | GTA | no | Clarion |
| Wake County | GTA | no | Clarion |
| Wake County | GTA | no | Clarion |
| Wake County | GTA | no | Clarion |
| Wake County | GTA | no | Clarion |
| Wake County | GTA | no | Clarion |
| Wake County | GTA | no | Clarion |
| Wake County | GTA | no | Clarion |
| Wake County | GTA | no | Clarion |
| Wake County | GTA | no | Clarion |
| Wake County | GTA | no | Clarion |
| Wake County | GTA | no | Clarion |
| Wake County | GTA | no | Clarion |
| Wake County | GTA | no | Clarion |
| Wake County | GTA | no | Clarion |
| Wake County | GTA | no | Clarion |
| Wake County | GTA | no | Clarion |
| Wake County | GTA | no | Clarion |
| Wake County | GTA | no | Clarion |
| Wake County | GTA | no | Clarion |

| Location | Position 1016 | KDR Mutant? | Site |
|---|---|---|---|
| Wake County | GTA | no | Clarion |
| Wake County | GTA | no | Clarion |
| Wake County | GTA | no | Clarion |
| Wake County | GTA | no | Clarion |
| Wake County | GTA | no | Clarion |
| Wake County | GTA | no | Clarion |
| Wake County | GTA | no | Clarion |
| Wake County | GTA | no | Clarion |
| Wake County | GTA | no | Clarion |
| Wake County | GTA | no | Clarion |
| Wake County | GTA | no | Clarion |
| Wake County | GTA | no | Clarion |
| Wake County | GTA | no | Clarion |
| Wake County | GTA | no | Clarion |
| Wake County | GTA | no | Clarion |
| Wake County | GTA | no | Clarion |
| Wake County | GTA | no | Clarion |
| Wake County | GTA | no | Clarion |
| Wake County | GTA | no | DIX |
| Wake County | GTA | no | DIX |
| Wake County | GTA | no | Farm |
| Wake County | GTA | no | Farm |
| Wake County | GTA | no | Farm |
| Wake County | GTA | no | Farm |
| Wake County | GTA | no | Farm |
| Wake County | GTA | no | Farm |

| Location | Position 1016 | KDR Mutant? | Site |
|---|---|---|---|
| Wake County | GTA | no | Farm |
| Wake County | GTA | no | Farm |
| Wake County | GTA | no | Farm |
| Wake County | GTA | no | Farm |
| Wake County | GTA | no | Farm |
| Wake County | GTA | no | Farm |
| Wake County | GTA | no | Farm |
| Wake County | GTA | no | Farm |
| Wake County | GTA | no | Farm |
| Wake County | GTA | no | Farm |
| Wake County | GTA | no | Farm |
| Wake County | GTA | no | Dairy |
| Wake County | GTA | no | Dairy |
| Wake County | GTA | no | Dairy |
| Wake County | GTA | no | Dairy |
| Wake County | GTA | no | Dairy |
| Wake County | GTA | no | Dairy |
| Wake County | GTA | no | Dairy |
| Wake County | GTA | no | Dairy |
| Wake County | GTA | no | Dairy |
| Wake County | GTA | no | Dairy |
| Wake County | GTA | no | Dairy |
| Wake County | GTA | no | Dairy |
| Wake County | GTA | no | Dairy |
| Wake County | GTA | no | Dairy |
| Wake County | GTA | no | Dairy |

| Location | Position 1016 | KDR Mutant? | Site |
|---|---|---|---|
| Wake County | GTA | no | Dairy |
| Wake County | GTA | no | Dairy |
| Wake County | GTA | no | Dairy |
| Wake County | GTA | no | Dairy |
| Wake County | GTA | no | Dairy |
| Wake County | GTA | no | Dairy |
| Wake County | GTA | no | Dairy |
| Wake County | GTA | no | Dairy |
| Wake County | GTA | no | Dairy |
| Wake County | GTA | no | Dairy |
| Wake County | GTA | no | Dairy |
| Wake County | GTA | no | Dairy |
| Wake County | GTA | no | Dairy |
| Wake County | GTA | no | Dairy |
| Wake County | GTA | no | Dairy |
| Wake County | GTA | no | Dairy |
| Wake County | GTA | no | Dairy |
| Wake County | GTA | no | Dairy |
| Wake County | GTA | no | Dairy |
| Wake County | GTA | no | Dairy |
| Wake County | GTA | no | Dairy |
| Wake County | GTA | no | Dairy |
| Wake County | GTA | no | Dairy |
| Wake County | GTA | no | Dairy |
| Wake County | GTA | no | Dairy |
| Wake County | GTA | no | Dairy |

| Location | Position 1016 | KDR Mutant? | Site |
|---|---|---|---|
| Wake County | GTA | no | Dairy |
| Wake County | GTA | no | Dairy |
| Wake County | GTA | no | Dairy |
| Wake County | GTA | no | Dairy |
| Wake County | GTA | no | Dairy |
| Wake County | GTA | no | Dairy |
| Wake County | GTA | no | Dairy |
| Wake County | GTA | no | Dairy |
| Wake County | GTA | no | Dairy |
| Wake County | GTA | no | Dairy |
| Wake County | GTA | no | Dairy |
| Wake County | GTA | no | Dairy |
| Wake County | GTA | no | Dairy |
| Wake County | GTA | no | Dairy |
| Wake County | GTA | no | Dairy |
| Wake County | GTA | no | Dairy |
| Wake County | GTA | no | Dairy |
| Wake County | GTA | no | Dairy |
| Wake County | GTA | no | Dairy |
| Wake County | GTA | no | Dairy |
| Wake County | GTA | no | Dairy |
| Wake County | GTA | no | Dairy |
| Wake County | GTA | no | Dairy |
| Wake County | GTA | no | Dairy |
| Wake County | GTA | no | Dairy |
| Wake County | GTA | no | Dairy |
| Wake County | GTA | no | Dairy |

| Location | Position 1016 | KDR Mutant? | Site |
|---|---|---|---|
| Wake County | GTA | no | Dairy |
| Wake County | GTA | no | Dairy |
| Wake County | GTA | no | Dairy |
| Wake County | GTA | no | Dairy |
| Wake County | GTA | no | Dairy |
| Wake County | GTA | no | Dairy |
| Wake County | GTA | no | Dairy |
| Wake County | GTA | no | Dairy |
| Wake County | GTA | no | Dairy |
| Wake County | GTA | no | Dairy |
| Wake County | GTA | no | Dairy |
| Wake County | GTA | no | Dairy |
| Wake County | GTA | no | BRF |
| Wake County | GTA | no | BRF |
| Wake County | GTA | no | BRF |
| Wake County | GTA | no | BRF |
| Wake County | GTA | no | BRF |
| Wake County | GTA | no | BRF |
| Wake County | GTA | no | BRF |
| Wake County | GTA | no | BRF |
| Wake County | GTA | no | BRF |
| Wake County | GTA | no | BRF |
| Wake County | GTA | no | BRF |
| Wake County | GTA | no | BRF |
| Wake County | GTA | no | BRF |
| Wake County | GTA | no | BRF |

| Location | Position 1016 | KDR Mutant? | Site |
|---|---|---|---|
| Wake County | GTA | no | BRF |
| Wake County | GTA | no | BRF |
| Wake County | GTA | no | BRF |
| Wake County | GTA | no | BRF |
| Wake County | GTA | no | BRF |
| Wake County | GTA | no | BRF |
| Wake County | GTA | no | BRF |
| Wake County | GTA | no | BRF |
| Wake County | GTA | no | BRF |
| Wake County | GTA | no | BRF |
| Wake County | GTA | no | BRF |
| Wake County | GTA | no | BRF |
| Wake County | GTA | no | BRF |
| Wake County | GTA | no | BRF |
| Wake County | GTA | no | BRF |
| Wake County | GTA | no | BRF |
| Wake County | GTA | no | BRF |
| Wake County | GTA | no | BRF |
| Wake County | GTA | no | BRF |
| Wake County | GTA | no | BRF |
| Wake County | GTA | no | BRF |
| Wake County | GTA | no | BRF |
| Wake County | GTA | no | BRF |
| Wake County | GTA | no | BRF |
| Wake County | GTA | no | BRF |
| Wake County | GTA | no | BRF |

| Location | Position 1016 | KDR Mutant? | Site |
|---|---|---|---|
| Wake County | GTA | no | BRF |
| Wake County | GTA | no | BRF |
| Wake County | GTA | no | BRF |
| Wake County | GTA | no | BRF |
| Wake County | GTA | no | BRF |
| Wake County | GTA | no | BRF |
| Wake County | GTA | no | BRF |
| Wake County | GTA | no | BRF |
| Wake County | GTA | no | VDR |
| Wake County | GTA | no | VDR |
| Wake County | GTA | no | VDR |
| Wake County | GTA | no | VDR |
| Wake County | GTA | no | VDR |
| Wake County | GTA | no | VDR |
| Wake County | GTA | no | VDR |
| Wake County | GTA | no | VDR |
| Wake County | GTA | no | VDR |
| Wake County | GTA | no | VDR |
| Wake County | GTA | no | VDR |
| Wake County | GTA | no | VDR |
| Wake County | GTA | no | VDR |
| Wake County | GTA | no | VDR |
| Wake County | GTA | no | VDR |
| Wake County | GTA | no | VDR |
| Wake County | GTA | no | VDR |
| Wake County | GTA | no | VDR |

| Location | Position 1016 | KDR Mutant? | Site |
|---|---|---|---|
| Wake County | GTA | no | VDR |
| Wake County | GTA | no | VDR |
| Wake County | GTA | no | VDR |
| Wake County | GTA | no | VDR |
| Wake County | GTA | no | VDR |
| Wake County | GTA | no | VDR |
| Wake County | GTA | no | VDR |
| Wake County | GTA | no | VDR |
| Wake County | GTA | no | VDR |
| Wake County | GTA | no | VDR |
| Wake County | GTA | no | VDR |
| Wake County | GTA | no | VDR |
| Wake County | GTA | no | VDR |
| Wake County | GTA | no | VDR |
| Wake County | GTA | no | VDR |
| Wake County | GTA | no | VDR |
| Wake County | GTA | no | VDR |
| Wake County | GTA | no | VDR |
| Wake County | GTA | no | VDR |
| Wake County | GTA | no | VDR |
| Wake County | GTA | no | Schenck |
| Wake County | GTA | no | Schenck |
| Wake County | GTA | no | MET |
| Wake County | GTA | no | MET |
| Wake County | GTA | no | MET |
| Wake County | GTA | no | MET |

| Location | Position 1016 | KDR Mutant? | Site |
|---|---|---|---|
| Wake County | GTA | no | MET |
| Wake County | GTA | no | MET |
| Wake County | GTA | no | Trexler |
| Wake County | GTA | no | Trexler |
| Wake County | GTA | no | Trexler |
| Wake County | GTA | no | Trexler |
| Wake County | GTA | no | Trexler |
| Wake County | GTA | no | Trexler |
| Wake County | GTA | no | Trexler |
| Wake County | GTA | no | Trexler |
| Wake County | GTA | no | Trexler |
| Wake County | GTA | no | Trexler |
| Wake County | GTA | no | Trexler |
| Wake County | GTA | no | Trexler |
| Wake County | GTA | no | Trexler |
| Wake County | GTA | no | Trexler |
| Wake County | GTA | no | Trexler |
| Wake County | GTA | no | Trexler |
| Wake County | GTA | no | Trexler |
| Wake County | GTA | no | Trexler |
| Wake County | GTA | no | Trexler |
| Wake County | GTA | no | Trexler |
| Wake County | GTA | no | Trexler |
| Wake County | GTA | no | Trexler |
| Wake County | GTA | no | Trexler |
| Wake County | GTA | no | Trexler |

| Location | Position 1016 | KDR Mutant? | Site |
| --- | --- | --- | --- |
| Wake County | GTA | no | Trexler |

**Table S4.** Domain three sequence data for each mosquito successfully sequenced.

| Location | Position 1532 | KDR Mutant? | Position 1534 | KDR Mutant? | Site |
|---|---|---|---|---|---|
| Wake County | ATC | No | TCC | yes | BRF |
| Wake County | ATC | No | TCC | yes | VDR |
| Wake County | ATC | No | TCC | yes | VDR |
| Wake County | ATC | No | TCC | yes | VDR |
| Wake County | ATC | No | TCC | yes | VDR |
| Wake County | ATC | No | TCC | yes | VDR |
| Wake County | ATC | No | TCC | yes | VDR |
| Wake County | ATC | No | TCC | yes | VDR |
| Wake County | ATC | No | TCC | yes | VDR |
| Wake County | ATC | No | TCC | yes | VDR |
| Wake County | ATC | No | TCC | yes | VDR |
| Wake County | ATC | No | TCC | yes | VDR |
| Fort Bragg | ATC | No | TTC | no | Woodland Heights |
| Fort Bragg | ATC | No | TTC | no | Woodland Heights |
| Fort Bragg | ATC | No | TTC | no | Woodland Heights |
| Fort Bragg | ATC | No | TTC | no | Woodland Heights |
| Fort Bragg | ATC | No | TTC | no | Woodland Heights |
| Fort Bragg | ATC | No | TTC | no | Woodland Heights |
| Fort Bragg | ATC | No | TTC | no | Woodland Heights |
| Fort Bragg | ATC | No | TTC | no | Woodland Heights |
| Fort Bragg | ATC | No | TTC | no | Woodland Heights |
| Fort Bragg | ATC | No | TTC | no | Woodland Heights |
| Fort Bragg | ATC | No | TTC | no | Woodland Heights |
| Fort Bragg | ATC | No | TTC | no | Woodland Heights |
| Fort Bragg | ATC | No | TTC | no | Woodland Heights |

| Location | Position 1532 | KDR Mutant? | Position 1534 | KDR Mutant? | Site |
|---|---|---|---|---|---|
| Fort Bragg | ATC | No | TTC | no | Ryder Golf Course |
| Fort Bragg | ATC | No | TTC | no | Ryder Golf Course |
| Fort Bragg | ATC | No | TTC | no | Ryder Golf Course |
| Fort Bragg | ATC | No | TTC | no | Ryder Golf Course |
| Fort Bragg | ATC | No | TTC | no | Ryder Golf Course |
| Fort Bragg | ATC | No | TTC | no | Nijmegen |
| Fort Bragg | ATC | No | TTC | no | Woodland Heights |
| Fort Bragg | ATC | No | TTC | no | Woodland Heights |
| Fort Bragg | ATC | No | TTC | no | Woodland Heights |
| Fort Bragg | ATC | No | TTC | no | Woodland Heights |
| Fort Bragg | ATC | No | TTC | no | Woodland Heights |
| Fort Bragg | ATC | No | TTC | no | Woodland Heights |
| Fort Bragg | ATC | No | TTC | no | Woodland Heights |
| Fort Bragg | ATC | No | TTC | no | Woodland Heights |
| Fort Bragg | ATC | No | TTC | no | Woodland Heights |
| Fort Bragg | ATC | No | TTC | no | Woodland Heights |
| Fort Bragg | ATC | No | TTC | no | Woodland Heights |
| Fort Bragg | ATC | No | TTC | no | Woodland Heights |
| Fort Bragg | ATC | No | TTC | no | Woodland Heights |
| Fort Bragg | ATC | No | TTC | no | Woodland Heights |
| Fort Bragg | ATC | No | TTC | no | Woodland Heights |
| Fort Bragg | ATC | No | TTC | no | Woodland Heights |
| Fort Bragg | ATC | No | TTC | no | Woodland Heights |
| Fort Bragg | ATC | No | TTC | no | Woodland Heights |
| Fort Bragg | ATC | No | TTC | no | Nijmegen |
| Fort Bragg | ATC | No | TTC | no | Nijmegen |

| Location | Position 1532 | KDR Mutant? | Position 1534 | KDR Mutant? | Site |
|---|---|---|---|---|---|
| Fort Bragg | ATC | No | TTC | no | Nijmegen |
| Fort Bragg | ATC | No | TTC | no | Nijmegen |
| Fort Bragg | ATC | No | TTC | no | Nijmegen |
| Fort Bragg | ATC | No | TTC | no | Nijmegen |
| Fort Bragg | ATC | No | TTC | no | Nijmegen |
| Fort Bragg | ATC | No | TTC | no | Nijmegen |
| Fort Bragg | ATC | No | TTC | no | Nijmegen |
| Fort Bragg | ATC | No | TTC | no | Nijmegen |
| Fort Bragg | ATC | No | TTC | no | Nijmegen |
| Fort Bragg | ATC | No | TTC | no | Nijmegen |
| Fort Bragg | ATC | No | TTC | no | Nijmegen |
| Fort Bragg | ATC | No | TTC | no | Nijmegen |
| Fort Bragg | ATC | No | TTC | no | Nijmegen |
| Fort Bragg | ATC | No | TTC | no | Nijmegen |
| Fort Bragg | ATC | No | TTC | no | Nijmegen |
| Fort Bragg | ATC | No | TTC | no | Nijmegen |
| Fort Bragg | ATC | No | TTC | no | Nijmegen |
| Fort Bragg | ATC | No | TTC | no | Nijmegen |
| Fort Bragg | ATC | No | TTC | no | Nijmegen |
| Fort Bragg | ATC | No | TTC | no | Ryder Golf Course |
| Fort Bragg | ATC | No | TTC | no | Ritz-Epps |
| Fort Bragg | ATC | No | TTC | no | Ritz-Epps |
| Fort Bragg | ATC | No | TTC | no | Ryder Golf Course |
| Fort Bragg | ATC | No | TTC | no | Ryder Golf Course |
| Fort Bragg | ATC | No | TTC | no | Ryder Golf Course |
| Fort Bragg | ATC | No | TTC | no | Ryder Golf Course |

| Location | Position 1532 | KDR Mutant? | Position 1534 | KDR Mutant? | Site |
|---|---|---|---|---|---|
| Fort Bragg | ATC | No | TTC | no | Ryder Golf Course |
| Fort Bragg | ATC | No | TTC | no | Nijmegen |
| Fort Bragg | ATC | No | TTC | no | Nijmegen |
| Fort Bragg | ATC | No | TTC | no | Nijmegen |
| Fort Bragg | ATC | No | TTC | no | Nijmegen |
| Fort Bragg | ATC | No | TTC | no | Nijmegen |
| Fort Bragg | ATC | No | TTC | no | Nijmegen |
| Fort Bragg | ATC | No | TTC | no | Nijmegen |
| Fort Bragg | ATC | No | TTC | no | Nijmegen |
| Fort Bragg | ATC | No | TTC | no | Nijmegen |
| Fort Bragg | ATC | No | TTC | no | Nijmegen |
| Fort Bragg | ATC | No | TTC | no | Nijmegen |
| Fort Bragg | ATC | No | TTC | no | Nijmegen |
| Fort Bragg | ATC | No | TTC | no | Nijmegen |
| Fort Bragg | ATC | No | TTC | no | Nijmegen |
| Fort Bragg | ATC | No | TTC | no | Nijmegen |
| Fort Bragg | ATC | No | TTC | no | Nijmegen |
| Fort Bragg | ATC | No | TTC | no | Ritz-Epps |
| Fort Bragg | ATC | No | TTC | no | Ritz-Epps |
| Fort Bragg | ATC | No | TTC | no | Woodland Heights |
| Fort Bragg | ATC | No | TTC | no | Woodland Heights |
| Fort Bragg | ATC | No | TTC | no | Woodland Heights |
| Fort Bragg | ATC | No | TTC | no | Woodland Heights |
| Fort Bragg | ATC | No | TTC | no | Woodland Heights |
| Fort Bragg | ATC | No | TTC | no | Woodland Heights |
| Fort Bragg | ATC | No | TTC | no | Woodland Heights |

| Location | Position 1532 | KDR Mutant? | Position 1534 | KDR Mutant? | Site |
|---|---|---|---|---|---|
| Fort Bragg | ATC | No | TTC | no | Woodland Heights |
| Fort Bragg | ATC | No | TTC | no | Woodland Heights |
| Fort Bragg | ATC | No | TTC | no | Woodland Heights |
| Fort Bragg | ATC | No | TTC | no | Woodland Heights |
| Fort Bragg | ATC | No | TTC | no | Woodland Heights |
| Fort Bragg | ATC | No | TTC | no | Woodland Heights |
| Fort Bragg | ATC | No | TTC | no | Woodland Heights |
| Fort Bragg | ATC | No | TTC | no | Ryder Golf Course |
| Fort Bragg | ATC | No | TTC | no | Ryder Golf Course |
| Fort Bragg | ATC | No | TTC | no | Ryder Golf Course |
| Fort Bragg | ATC | No | TTC | no | Ryder Golf Course |
| Fort Bragg | ATC | No | TTC | no | Ryder Golf Course |
| Fort Bragg | ATC | No | TTC | no | Ryder Golf Course |
| Fort Bragg | ATC | No | TTC | no | Ryder Golf Course |
| Fort Bragg | ATC | No | TTC | no | Ryder Golf Course |
| Fort Bragg | ATC | No | TTC | no | Ryder Golf Course |
| Fort Bragg | ATC | No | TTC | no | Ryder Golf Course |
| Fort Bragg | ATC | No | TTC | no | Ryder Golf Course |
| Fort Bragg | ATC | No | TTC | no | Ryder Golf Course |
| Fort Bragg | ATC | No | TTC | no | Ryder Golf Course |
| Fort Bragg | ATC | No | TTC | no | Nijmegen |
| Fort Bragg | ATC | No | TTC | no | Nijmegen |
| Fort Bragg | ATC | No | TTC | no | Nijmegen |
| Fort Bragg | ATC | No | TTC | no | Nijmegen |
| Fort Bragg | ATC | No | TTC | no | Nijmegen |

| Location | Position 1532 | KDR Mutant? | Position 1534 | KDR Mutant? | Site |
|---|---|---|---|---|---|
| Fort Bragg | ATC | No | TTC | no | Nijmegen |
| Fort Bragg | ATC | No | TTC | no | Nijmegen |
| Fort Bragg | ATC | No | TTC | no | Nijmegen |
| Fort Bragg | ATC | No | TTC | no | Nijmegen |
| Fort Bragg | ATC | No | TTC | no | Nijmegen |
| Fort Bragg | ATC | No | TTC | no | Nijmegen |
| Fort Bragg | ATC | No | TTC | no | Nijmegen |
| Fort Bragg | ATC | No | TTC | no | Nijmegen |
| Fort Bragg | ATC | No | TTC | no | Nijmegen |
| Fort Bragg | ATC | No | TTC | no | Nijmegen |
| Fort Bragg | ATC | No | TTC | no | Nijmegen |
| Fort Bragg | ATC | No | TTC | no | Nijmegen |
| Fort Bragg | ATC | No | TTC | no | Nijmegen |
| Fort Bragg | ATC | No | TTC | no | Nijmegen |
| Fort Bragg | ATC | No | TTC | no | Nijmegen |
| Fort Bragg | ATC | No | TTC | no | Nijmegen |
| Fort Bragg | ATC | No | TTC | no | Nijmegen |
| Wake County | ATC | No | TTC | no | Mission |
| Wake County | ATC | No | TTC | no | Mission |
| Wake County | ATC | No | TTC | no | Mission |
| Wake County | ATC | No | TTC | no | Mission |
| Wake County | ATC | No | TTC | no | Mission |
| Wake County | ATC | No | TTC | no | Mission |
| Wake County | ATC | No | TTC | no | Mission |
| Wake County | ATC | No | TTC | no | Mission |
| Wake County | ATC | No | TTC | no | Mission |

| Location | Position 1532 | KDR Mutant? | Position 1534 | KDR Mutant? | Site |
|---|---|---|---|---|---|
| Wake County | ATC | No | TTC | no | Mission |
| Wake County | ATC | No | TTC | no | Mission |
| Wake County | ATC | No | TTC | no | Mission |
| Wake County | ATC | No | TTC | no | Mission |
| Wake County | ATC | No | TTC | no | Mission |
| Wake County | ATC | No | TTC | no | Mission |
| Wake County | ATC | No | TTC | no | Mission |
| Wake County | ATC | No | TTC | no | Mission |
| Wake County | ATC | No | TTC | no | Mission |
| Wake County | ATC | No | TTC | no | Mission |
| Wake County | ATC | No | TTC | no | Market |
| Wake County | ATC | No | TTC | no | Market |
| Wake County | ATC | No | TTC | no | Market |
| Wake County | ATC | No | TTC | no | Market |
| Wake County | ATC | No | TTC | no | Market |
| Wake County | ATC | No | TTC | no | Forensic |
| Wake County | ATC | No | TTC | no | Forensic |
| Wake County | ATC | No | TTC | no | Forensic |
| Wake County | ATC | No | TTC | no | Forensic |
| Wake County | ATC | No | TTC | no | Forensic |
| Wake County | ATC | No | TTC | no | Grinnells |
| Wake County | ATC | No | TTC | no | Grinnells |
| Wake County | ATC | No | TTC | no | Grinnells |
| Wake County | ATC | No | TTC | no | Varsity |
| Wake County | ATC | No | TTC | no | Varsity |
| Wake County | ATC | No | TTC | no | Varsity |

| Location | Position 1532 | KDR Mutant? | Position 1534 | KDR Mutant? | Site |
|---|---|---|---|---|---|
| Wake County | ATC | No | TTC | no | Varsity |
| Wake County | ATC | No | TTC | no | Varsity |
| Wake County | ATC | No | TTC | no | Varsity |
| Wake County | ATC | No | TTC | no | Varsity |
| Wake County | ATC | No | TTC | no | Varsity |
| Wake County | ATC | No | TTC | no | Varsity |
| Wake County | ATC | No | TTC | no | Varsity |
| Wake County | ATC | No | TTC | no | Varsity |
| Wake County | ATC | No | TTC | no | Clarion |
| Wake County | ATC | No | TTC | no | Clarion |
| Wake County | ATC | No | TTC | no | Clarion |
| Wake County | ATC | No | TTC | no | Clarion |
| Wake County | ATC | No | TTC | no | Clarion |
| Wake County | ATC | No | TTC | no | Clarion |
| Wake County | ATC | No | TTC | no | Clarion |
| Wake County | ATC | No | TTC | no | Clarion |
| Wake County | ATC | No | TTC | no | Clarion |
| Wake County | ATC | No | TTC | no | Clarion |
| Wake County | ATC | No | TTC | no | Clarion |
| Wake County | ATC | No | TTC | no | Clarion |
| Wake County | ATC | No | TTC | no | Clarion |
| Wake County | ATC | No | TTC | no | Clarion |
| Wake County | ATC | No | TTC | no | Clarion |
| Wake County | ATC | No | TTC | no | Clarion |
| Wake County | ATC | No | TTC | no | Clarion |
| Wake County | ATC | No | TTC | no | Clarion |

| Location | Position 1532 | KDR Mutant? | Position 1534 | KDR Mutant? | Site |
|---|---|---|---|---|---|
| Wake County | ATC | No | TTC | no | Clarion |
| Wake County | ATC | No | TTC | no | Clarion |
| Wake County | ATC | No | TTC | no | Clarion |
| Wake County | ATC | No | TTC | no | Clarion |
| Wake County | ATC | No | TTC | no | Clarion |
| Wake County | ATC | No | TTC | no | Clarion |
| Wake County | ATC | No | TTC | no | Clarion |
| Wake County | ATC | No | TTC | no | Clarion |
| Wake County | ATC | No | TTC | no | Clarion |
| Wake County | ATC | No | TTC | no | Clarion |
| Wake County | ATC | No | TTC | no | Clarion |
| Wake County | ATC | No | TTC | no | Clarion |
| Wake County | ATC | No | TTC | no | Clarion |
| Wake County | ATC | No | TTC | no | Clarion |
| Wake County | ATC | No | TTC | no | Clarion |
| Wake County | ATC | No | TTC | no | Clarion |
| Wake County | ATC | No | TTC | no | Clarion |
| Wake County | ATC | No | TTC | no | Clarion |
| Wake County | ATC | No | TTC | no | Clarion |
| Wake County | ATC | No | TTC | no | Clarion |
| Wake County | ATC | No | TTC | no | DIX |
| Wake County | ATC | No | TTC | no | DIX |
| Wake County | ATC | No | TTC | no | Farm |
| Wake County | ATC | No | TTC | no | Farm |
| Wake County | ATC | No | TTC | no | Farm |
| Wake County | ATC | No | TTC | no | Farm |

| Location | Position 1532 | KDR Mutant? | Position 1534 | KDR Mutant? | Site |
|---|---|---|---|---|---|
| Wake County | ATC | No | TTC | no | Farm |
| Wake County | ATC | No | TTC | no | Farm |
| Wake County | ATC | No | TTC | no | Farm |
| Wake County | ATC | No | TTC | no | Farm |
| Wake County | ATC | No | TTC | no | Farm |
| Wake County | ATC | No | TTC | no | Farm |
| Wake County | ATC | No | TTC | no | Farm |
| Wake County | ATC | No | TTC | no | Farm |
| Wake County | ATC | No | TTC | no | Farm |
| Wake County | ATC | No | TTC | no | Farm |
| Wake County | ATC | No | TTC | no | Farm |
| Wake County | ATC | No | TTC | no | Farm |
| Wake County | ATC | No | TTC | no | Farm |
| Wake County | ATC | No | TTC | no | Farm |
| Wake County | ATC | No | TTC | no | Dairy |
| Wake County | ATC | No | TTC | no | Dairy |
| Wake County | ATC | No | TTC | no | Dairy |
| Wake County | ATC | No | TTC | no | Dairy |
| Wake County | ATC | No | TTC | no | Dairy |
| Wake County | ATC | No | TTC | no | Dairy |
| Wake County | ATC | No | TTC | no | Dairy |
| Wake County | ATC | No | TTC | no | Dairy |
| Wake County | ATC | No | TTC | no | Dairy |
| Wake County | ATC | No | TTC | no | Dairy |
| Wake County | ATC | No | TTC | no | Dairy |
| Wake County | ATC | No | TTC | no | Dairy |

| Location | Position 1532 | KDR Mutant? | Position 1534 | KDR Mutant? | Site |
|---|---|---|---|---|---|
| Wake County | ATC | No | TTC | no | Dairy |
| Wake County | ATC | No | TTC | no | Dairy |
| Wake County | ATC | No | TTC | no | Dairy |
| Wake County | ATC | No | TTC | no | Dairy |
| Wake County | ATC | No | TTC | no | Dairy |
| Wake County | ATC | No | TTC | no | Dairy |
| Wake County | ATC | No | TTC | no | Dairy |
| Wake County | ATC | No | TTC | no | Dairy |
| Wake County | ATC | No | TTC | no | Dairy |
| Wake County | ATC | No | TTC | no | Dairy |
| Wake County | ATC | No | TTC | no | Dairy |
| Wake County | ATC | No | TTC | no | Dairy |
| Wake County | ATC | No | TTC | no | Dairy |
| Wake County | ATC | No | TTC | no | Dairy |
| Wake County | ATC | No | TTC | no | Dairy |
| Wake County | ATC | No | TTC | no | Dairy |
| Wake County | ATC | No | TTC | no | Dairy |
| Wake County | ATC | No | TTC | no | Dairy |
| Wake County | ATC | No | TTC | no | Dairy |
| Wake County | ATC | No | TTC | no | Dairy |
| Wake County | ATC | No | TTC | no | Dairy |
| Wake County | ATC | No | TTC | no | Dairy |
| Wake County | ATC | No | TTC | no | Dairy |
| Wake County | ATC | No | TTC | no | Dairy |
| Wake County | ATC | No | TTC | no | Dairy |
| Wake County | ATC | No | TTC | no | Dairy |

| Location | Position 1532 | KDR Mutant? | Position 1534 | KDR Mutant? | Site |
|---|---|---|---|---|---|
| Wake County | ATC | No | TTC | no | Dairy |
| Wake County | ATC | No | TTC | no | Dairy |
| Wake County | ATC | No | TTC | no | Dairy |
| Wake County | ATC | No | TTC | no | Dairy |
| Wake County | ATC | No | TTC | no | Dairy |
| Wake County | ATC | No | TTC | no | Dairy |
| Wake County | ATC | No | TTC | no | Dairy |
| Wake County | ATC | No | TTC | no | Dairy |
| Wake County | ATC | No | TTC | no | Dairy |
| Wake County | ATC | No | TTC | no | Dairy |
| Wake County | ATC | No | TTC | no | Dairy |
| Wake County | ATC | No | TTC | no | Dairy |
| Wake County | ATC | No | TTC | no | Dairy |
| Wake County | ATC | No | TTC | no | Dairy |
| Wake County | ATC | No | TTC | no | Dairy |
| Wake County | ATC | No | TTC | no | Dairy |
| Wake County | ATC | No | TTC | no | Dairy |
| Wake County | ATC | No | TTC | no | Dairy |
| Wake County | ATC | No | TTC | no | Dairy |
| Wake County | ATC | No | TTC | no | Dairy |
| Wake County | ATC | No | TTC | no | Dairy |
| Wake County | ATC | No | TTC | no | Dairy |
| Wake County | ATC | No | TTC | no | Dairy |
| Wake County | ATC | No | TTC | no | Dairy |
| Wake County | ATC | No | TTC | no | Dairy |
| Wake County | ATC | No | TTC | no | Dairy |

| Location | Position 1532 | KDR Mutant? | Position 1534 | KDR Mutant? | Site |
|---|---|---|---|---|---|
| Wake County | ATC | No | TTC | no | Dairy |
| Wake County | ATC | No | TTC | no | Dairy |
| Wake County | ATC | No | TTC | no | Dairy |
| Wake County | ATC | No | TTC | no | Dairy |
| Wake County | ATC | No | TTC | no | Dairy |
| Wake County | ATC | No | TTC | no | BRF |
| Wake County | ATC | No | TTC | no | BRF |
| Wake County | ATC | No | TTC | no | BRF |
| Wake County | ATC | No | TTC | no | BRF |
| Wake County | ATC | No | TTC | no | BRF |
| Wake County | ATC | No | TTC | no | BRF |
| Wake County | ATC | No | TTC | no | BRF |
| Wake County | ATC | No | TTC | no | BRF |
| Wake County | ATC | No | TTC | no | BRF |
| Wake County | ATC | No | TTC | no | BRF |
| Wake County | ATC | No | TTC | no | BRF |
| Wake County | ATC | No | TTC | no | BRF |
| Wake County | ATC | No | TTC | no | BRF |
| Wake County | ATC | No | TTC | no | BRF |
| Wake County | ATC | No | TTC | no | BRF |
| Wake County | ATC | No | TTC | no | BRF |
| Wake County | ATC | No | TTC | no | BRF |
| Wake County | ATC | No | TTC | no | BRF |
| Wake County | ATC | No | TTC | no | BRF |
| Wake County | ATC | No | TTC | no | BRF |
| Wake County | ATC | No | TTC | no | BRF |

| Location | Position 1532 | KDR Mutant? | Position 1534 | KDR Mutant? | Site |
|---|---|---|---|---|---|
| Wake County | ATC | No | TTC | no | BRF |
| Wake County | ATC | No | TTC | no | BRF |
| Wake County | ATC | No | TTC | no | BRF |
| Wake County | ATC | No | TTC | no | BRF |
| Wake County | ATC | No | TTC | no | BRF |
| Wake County | ATC | No | TTC | no | BRF |
| Wake County | ATC | No | TTC | no | BRF |
| Wake County | ATC | No | TTC | no | BRF |
| Wake County | ATC | No | TTC | no | BRF |
| Wake County | ATC | No | TTC | no | VDR |
| Wake County | ATC | No | TTC | no | VDR |
| Wake County | ATC | No | TTC | no | VDR |
| Wake County | ATC | No | TTC | no | VDR |
| Wake County | ATC | No | TTC | no | VDR |
| Wake County | ATC | No | TTC | no | VDR |
| Wake County | ATC | No | TTC | no | VDR |
| Wake County | ATC | No | TTC | no | VDR |
| Wake County | ATC | No | TTC | no | VDR |
| Wake County | ATC | No | TTC | no | VDR |
| Wake County | ATC | No | TTC | no | VDR |
| Wake County | ATC | No | TTC | no | VDR |
| Wake County | ATC | No | TTC | no | VDR |
| Wake County | ATC | No | TTC | no | VDR |
| Wake County | ATC | No | TTC | no | VDR |
| Wake County | ATC | No | TTC | no | VDR |
| Wake County | ATC | No | TTC | no | VDR |

| Location | Position 1532 | KDR Mutant? | Position 1534 | KDR Mutant? | Site |
|---|---|---|---|---|---|
| Wake County | ATC | No | TTC | no | VDR |
| Wake County | ATC | No | TTC | no | VDR |
| Wake County | ATC | No | TTC | no | VDR |
| Wake County | ATC | No | TTC | no | VDR |
| Wake County | ATC | No | TTC | no | VDR |
| Wake County | ATC | No | TTC | no | VDR |
| Wake County | ATC | No | TTC | no | VDR |
| Wake County | ATC | No | TTC | no | VDR |
| Wake County | ATC | No | TTC | no | VDR |
| Wake County | ATC | No | TTC | no | VDR |
| Wake County | ATC | No | TTC | no | VDR |
| Wake County | ATC | No | TTC | no | VDR |
| Wake County | ATC | No | TTC | no | VDR |
| Wake County | ATC | No | TTC | no | VDR |
| Wake County | ATC | No | TTC | no | VDR |
| Wake County | ATC | No | TTC | no | VDR |
| Wake County | ATC | No | TTC | no | VDR |
| Wake County | ATC | No | TTC | no | Schenck |
| Wake County | ATC | No | TTC | no | Schenck |
| Wake County | ATC | No | TTC | no | MET |
| Wake County | ATC | No | TTC | no | MET |
| Wake County | ATC | No | TTC | no | MET |
| Wake County | ATC | No | TTC | no | MET |
| Wake County | ATC | No | TTC | no | MET |
| Wake County | ATC | No | TTC | no | MET |
| Wake County | ATC | No | TTC | no | Trexler |

| Location | Position 1532 | KDR Mutant? | Position 1534 | KDR Mutant? | Site |
|---|---|---|---|---|---|
| Wake County | ATC | No | TTC | no | Trexler |
| Wake County | ATC | No | TTC | no | Trexler |
| Wake County | ATC | No | TTC | no | Trexler |
| Wake County | ATC | No | TTC | no | Trexler |
| Wake County | ATC | No | TTC | no | Trexler |
| Wake County | ATC | No | TTC | no | Trexler |
| Wake County | ATC | No | TTC | no | Trexler |
| Wake County | ATC | No | TTC | no | Trexler |
| Wake County | ATC | No | TTC | no | Trexler |
| Wake County | ATC | No | TTC | no | Trexler |
| Wake County | ATC | No | TTC | no | Trexler |
| Wake County | ATC | No | TTC | no | Trexler |
| Wake County | ATC | No | TTC | no | Trexler |
| Wake County | ATC | No | TTC | no | Trexler |
| Wake County | ATC | No | TTC | no | Trexler |
| Wake County | ATC | No | TTC | no | Trexler |
| Wake County | ATC | No | TTC | no | Trexler |
| Wake County | ATC | No | TTC | no | Trexler |
| Wake County | ATC | No | TTC | no | Trexler |
| Wake County | ATC | No | TTC | no | Trexler |
| Wake County | ATC | No | TTC | no | Trexler |
| Wake County | ATC | No | TTC | no | Trexler |
| Wake County | ATC | No | TTC | no | Trexler |
